# A 5’ UTR G-quadruplex controls localisation and translation of a potassium leak channel mRNA

**DOI:** 10.1101/797423

**Authors:** Connor J. Maltby, James P. R. Schofield, Steven D. Houghton, Ita O’Kelly, Mariana Vargas-Caballero, Katrin Deinhardt, Mark J. Coldwell

## Abstract

RNA G-quadruplexes (G4s) are non-canonical secondary structures that have been proposed to function as regulators of post-transcriptional mRNA localisation and translation. G4s within 3’ UTRs of some neuronal mRNAs are known to control their distal localisation and local translation, contributing to the distinct local proteomes that facilitate the synaptic remodelling attributed to normal cellular function. In this study, we characterise the G4 formation of a (GGN)_13_ repeat found within the 5’ UTR of KCNK9 mRNA, encoding the potassium 2-pore domain leak channel Task3. Using circular dichroism, we show that this (GGN)_13_ repeat forms a parallel G4 that exhibits the stereotypical potassium specificity of a G4, remaining thermostable under physiological ionic conditions. The G4 is inhibitory to translation of Task3, which can be overcome through the activity of the G4-specific helicase DHX36, consequently increasing K^+^ leak currents and decreasing resting membrane potentials in HEK293 cells. Additionally, we observe that this G4 is fundamental to ensuring the delivery of Task3 mRNA to distal primary cortical neurites. It has previously been shown that abnormal Task3 expression correlates with neuronal dysfunction, we therefore posit that this G4 is required for regulated local expression of Task3 leak channels that maintain K^+^ leak currents within neurons.

## INTRODUCTION

DNA and RNA can adopt many secondary structures that have important roles in dictating cellular processes (1). G-quadruplexes (G4s) form in portions of guanine-rich DNA and RNA, and are amongst the most stable of nucleic acid secondary structures (2). G4s are more stable in RNA than DNA and can exist as mono- or multi-stranded structures (3). G4s compose of stacks of guanine tetrads that stabilise via non-canonical Hoogsteen hydrogen bonding, as opposed to the canonical Watson-Crick bonding observed in duplex DNA/RNA (Figure 1A). Stacked G-tetrads are stabilised through interactions with monovalent cations (4), where K^+^ elicits the greatest stabilisation whereas Li^+^ is unable to stabilise G4 formation due to constraints in ionic radii within the G4 central pore (5, 6).

**Figure 1.**
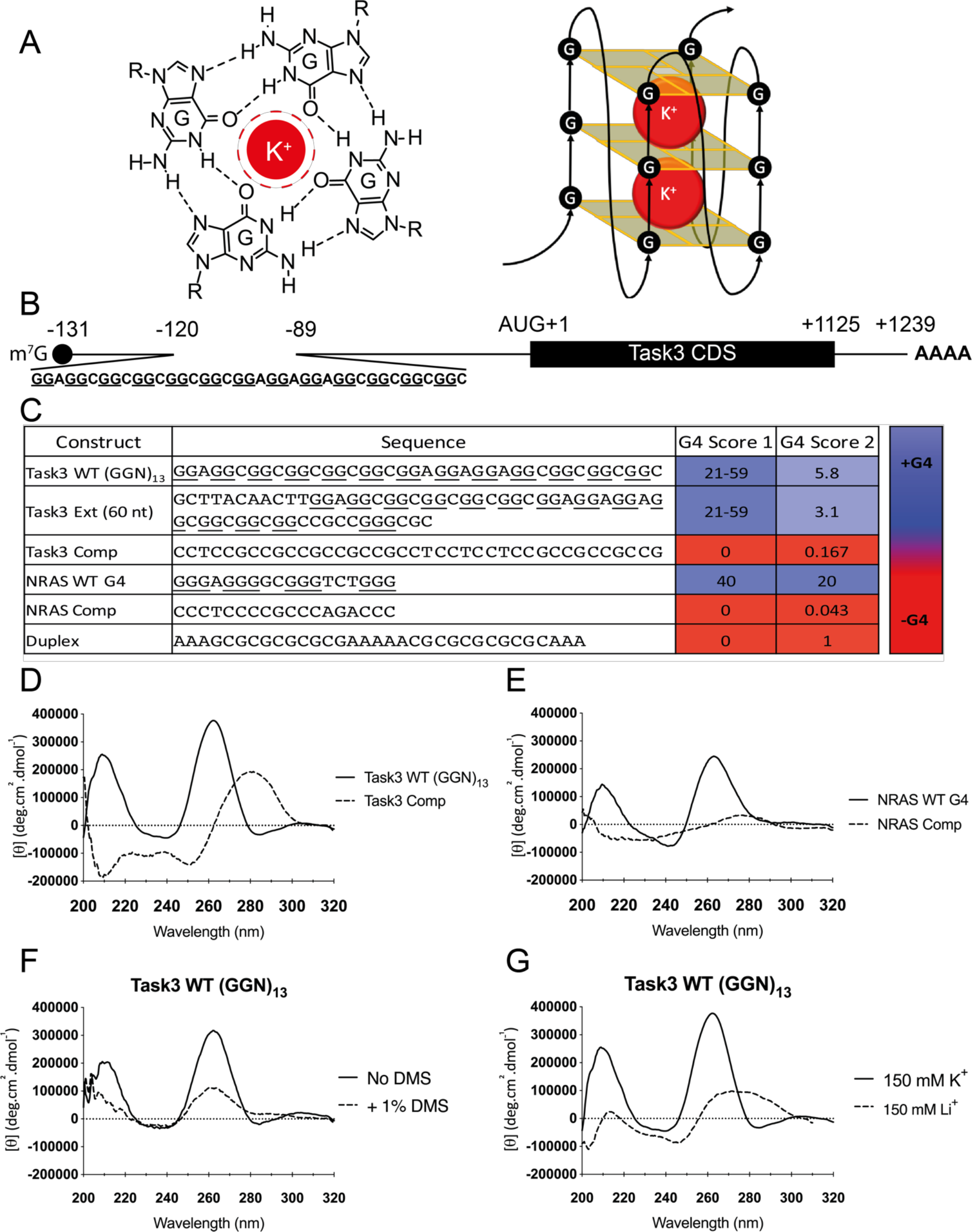
Task3 mRNA contains a 5’ UTR cap-proximal (GGN)_13_ repeat that forms a parallel G4. (**A**) Schematic illustrating conformation of Hoogsteen bonded guanine-quartets (left), that stack into G4 structures stabilised by K^+^ ions (right). (**B**) Map of Task3 mRNA indicating the (GGN)_13_ repeat within the 5’ UTR, as identified by 5’ RACE (GenBank: MN510330). (**C**) *In silico* prediction of Task3 experimental oligos structural formation based upon nucleotide composition and location – G4 score 1 and 2 refer to the G4 prediction tools outlined in methods. The CD spectra of 10 μM Task3 WT (GGN)_13_ and Task3 Complementary oligos (**D**) or NRAS WT G4 or NRAS Complementary oligos (**E**) folded in potassium phosphate buffer supplemented with KCl to 150 mM K^+^. (**F**) The CD spectra of 10 μM Task3 WT (GGN)_13_ oligos folded with or without pre-treatment of 1% dimethyl sulphate (DMS) to induce N7 methylation of guanine. (**G**) The CD spectra of 10 μM Task3 WT (GGN)_13_ oligos folded in potassium phosphate buffer supplemented with KCl to 150 mM K^+^ to facilitate G4 formation (CD spectra overlaid from 1D) or folded in sodium phosphate buffer supplemented with LiCl to 150 mM Li^+^ to inhibit G4 formation. CD spectra were obtained from five repeated acquisitions to increase the signal:noise ratio with a representative trace shown.

G4s are enriched within regulatory elements of both genomic DNA and mRNAs (7, 8), clustering at telomeres (9), promoter regions and transcription start sites (10–12) of genomic DNA, as well as splice sites (13) and untranslated regions of mRNAs (14–16). G4s in mRNAs are thought to be of particular importance in the regulation of gene expression, with G4s within the 5’ untranslated region (UTR) of mRNAs able to inhibit translation by impeding scanning of the 40S ribosomal subunit, or promoting translation through non-canonical ribosomal interactions with internal ribosomal entry sites (IRES) (17). Similarly, G4s within 3’ UTRs can impede translation through association with regulatory proteins (18), or indirectly through miRNA recruitment (19) as well as regulating polyadenylation (20) and subcellular localisation of mRNAs (21, 22).

Local translation is of particular importance within highly polarised cells, such as within neurons where singular cortical axons can project contra-laterally across the width of the brain. As such, both protein and mRNA transport to these distal sites of neurons is critical to neuronal homeostatic functions (23). Recently, nearly 2000 actively translating mRNAs were identified within the axon through an axon-specific translating ribosome affinity purification (TRAP) approach (24, 25). Similarly, identification of translating polyribosomes and over 2500 distinct mRNA transcripts through deep-sequencing within the hippocampal neuropil (26, 27) further suggests that neurons have a dependency on the local translation of mRNAs in distal neurite compartments (23, 28).

Disruption of G4 homeostasis has been implicated in several neurodevelopmental and degenerative diseases. A G4-forming (CGG)_n_ repeat expansion within mutant forms of the *FMR1* gene locus induces aberrant expression of the G4 binding protein FMRP which leads to the neurodevelopmental Fragile-X syndrome (FXS), or the neurodegenerative Fragile-X tremor and ataxia syndrome (FXTAS) (29). Similarly, the G4-forming hexanucleotide (GGGGCC)_n_ repeat expansion within the spliced intron 1 of C9Orf72 causes the most common form of inherited amyotrophic lateral sclerosis (ALS), and/or frontotemporal dementia (FTD), through RNA toxicity and repeat-associated non-AUG (RAN) translation producing toxic dipeptide-repeat species (30). As such, the importance of G4s during development and maintenance of synaptic connections through regulated mRNA localisation, as well as their role during neuronal dysfunction and neurodegeneration (29, 31, 32) is of particular interest.

The TWIK-related acid-sensitive K^+^ channel (Task3, also known as KCNK9 and K2P9.1) is a potassium two-pore domain (K2P) leak channel responsible for regulating the resting membrane potentials of various cell types (33). Task3 is enriched in brain regions associated with high rates of neuronal firing such as the hippocampus and cerebellum (34), and is required during processes such as synaptic plasticity. Task3 loss of function leads to the maternally-imprinted intellectual disability disorder Birk-Barel mental retardation (35), and perturbations in Task3 expression lead to changes in resting membrane potentials and neuronal action potential kinetics and recovery (34), suggesting that tight regulation of expression is essential. In this study, we analyse a novel (GGN)_13_ repeat in the cap-proximal region of the 5’ UTR of Task3 mRNA (Figure 1B) that folds into a parallel G4 structure responsible for the regulation of distal Task3 mRNA localisation and translational regulation required for the maintenance of normal cellular physiology.

## MATERIAL AND METHODS

### Materials

Primary antibodies used in this study were anti-FLAG M2 (Merck, F1804), anti-α-Actin (Sigma, AC-40), anti-Task3 (Merck, K0514), anti-DHX36 (Proteintech, 13159), anti-α-Tubulin (Proteintech, 66031), anti-Lamin (Cell Signalling Technology, 2032). Secondary antibodies used were conjugated to Alexa Fluor® fluorophores (Invitrogen) or IRDyes (LI-COR Biosciences). The plasmids used were cloned from human brain cDNA (Agilent 540005-41) into C-terminal pcDNA3.1-3F/eGFP vectors as previously described (36). Primers were obtained from Sigma, sequences of which are outlined in Figure S5.

Stellaris® RNA FISH probes were obtained from LGC Biosearch Tech. Anti-Task3 mouse RNA FISH probes were designed using the Stellaris® online web tool (see data availability) and conjugated to Quasar® 570 nm fluorophores. Anti-eGFP RNA FISH probes were pre-designed from Stellaris® and conjugated to Cal Fluor® Red 590 nm fluorophores.

### Circular Dichroism Spectroscopy

Oligonucleotides were dissolved in nuclease-free H_2_O to a concentration of 1 mM stock solution and diluted to an experimental concentration of 10 μM, estimated through OD_260_ absorption, in buffer containing potassium or sodium phosphate and potassium/lithium chloride salt to the desired free [K^+^/Li^+^]. The oligonucleotide preparations were then folded by heating to 95 °C for 10 mins and cooled at a rate of 0.5 °C/min to room temperature (RT).

CD spectra were recorded using a Jasco J-720 spectropolarimeter equipped with a Jasco PS-450 xenon lamp and a Grant LTD6G temperature-controlled cell holder. Quartz cuvettes of 1 mm path length were used and scans were recorded at a scan speed of 50 nm/min with response time of 2 s. The wavelength of scans was recorded between 320-200 nm, with 0.5 nm resolution and sensitivity of 20 m°. The average of 5 spectral scans was taken for each sample followed by baseline buffer correction. All scans were conducted at 25 °C except where temperature is the designated independent variable. The data were normalised to molar ellipticity using the equation: [θ] = (100 × θ)/(c × l), where c = molar concentration and l = pathlength in cm.

Temperature controlled experiments were achieved through water-conductive heating of the cell holder. Temperatures were increased from 20 °C to 95 °C at 5 °C increments, allowing 5 minutes stabilisation at each temperature before spectra were obtained. Samples were overlaid with mineral oil to prevent evaporation.

### Cell Culture

HEK293 cells were maintained in DMEM (Dulbecco’s Modified Eagle’s Medium) media (Gibco) with 10% FBS supplement (Gibco) and incubated at 37 °C and 5% CO_2_. Cells were transfected using Genejuice^TM^ (Novagen) as described (37) and proteins extracted after 48 h.

For tissue collection, C57Bl/6 wildtype mice were sacrificed in accordance with the Animals (Scientific Procedures) Act 1986 as approved by the UK Home Office. Primary neurons were prepared in Dulbecco’s PBS (DPBS, Life Technologies) from cortices of E15–E18 mice. Neurons were dissociated and maintained in complete NBM (Neurobasal medium with 2% (v/v) B27, 0.5 mM Glutamax) as previously described (38). The number of cells was determined and plated prior to transfection. For immunocytochemistry/RNA FISH, cells were transfected on day in vitro (DIV) 7 with Lipofectamine 2000 as described (39). The lipoplex mixture was added to the cells and incubated at 37 °C for 40 min. Subsequently, the lipoplex-containing medium was removed from the wells and replaced with conditioned medium, and the cells were returned to 37 °C. For bicuculline treatment, the cells were treated with 50 μM (+)-bicuculline (Tocris) or 0.1% DMSO (vehicle treatment) prepared in complete NBM. Cells were incubated at 37 °C for 24, 12, 6, 3, 1 h and 15 mins and then harvested. For glutamate treatment, the cells were treated with 50 μM L-glutamic acid, monosodium salt monohydrate (Sigma) dissolved in ddH_2_O prepared in complete NBM conjunction with a media change control. Cells were incubated for 30, 60 or 120 s, media changed and then allowed to recover for 1 h at 37 °C before being harvested.

For protein/RNA isolation using Corning^®^ Costar^®^ Transwell^®^ inserts (24 mm diameter, 3491), murine embryonic cortical neurons were seeded at the desired density and protein/RNA harvested at DIV7. Axonal preparations were isolated as described (40).

### HEK293 Cell Recording

HEK293 cells were maintained and transfected as above and whole-cell patch clamp was performed after 24 h transfection. Thick-glass borosilicate glass micropipettes were pulled to a tip resistance of 5-7 mOhm then filled with intracellular recording solution containing: 115 mM potassium gluconate, 10 mM KCl, 10 mM HEPES, 10 mM potassium phosphocreatine, 0.4 mM GTP, 4 mM ATP, and pH balanced to 7.3 with KOH. Osmolarity of ~280 mOsm. Cells were perfused with artificial cerebrospinal fluid (ACSF) kept at 37 °C throughout recording. ACSF contained: 126 mM NaCl, 2 mM CaCl_2_, 10 mM glucose, 2 mM MgSO_4_.7H_2_O, 3 mM KCl, 1.25 mM NaH_2_PO4.2H_2_O and 26.4 mM NaHCO_3_, bubbled with carbogen gas (95% O_2_, 5% CO_2_). Osmolarity of ~ 300 mOsm. The membrane potential was measured using a Multiclamp 700B (Molecular Devices) in current clamp (bridge) mode. Membrane potential was corrected for liquid junction potential of −12.5 mV which was measured directly (41).

### Western Blotting

Proteins were extracted in Radioimmunoprecipitation assay (RIPA) buffer (150 mM NaCl, 1% (v/v) NP-40, 1% (w/v) DOC, 0.1% (v/v) SDS, 50 mM Tris HCL pH 7.6, 1 mM EDTA, 1 mM EGTA, 1× Halt protease and phosphatase inhibitor cocktail (Thermo)) as previously described (42). Protein lysates were prepared and blotted following standard procedures, and images were captured using the Odyssey IR Imaging System (LI-COR Biosciences). Image Studio Scanner software was used to obtain the image, and Image Studio Lite software was used to quantify the intensities of the bands.

### RT-PCR and RT-qPCR

RNA was extracted using the NucleoSpin^®^ RNA extraction kit (Machery Nagel) according to manufacturer’s instructions. RNA was reverse transcribed to cDNA using the qScript cDNA Supermix (QuantaBio) according to manufacturer’s instructions. Endpoint PCRs were carried out using Readymix™ Taq PCR Reaction Mix (Sigma) with primers obtained from Sigma (see materials and Figure S5). Quantitative PCR (qPCR) amplification was performed using SYBR® Select Master Mix (Life Technologies) and an Eco qPCR System (Illumina) using the following conditions: initial denaturation 95 °C for 5 min, 40 cycles of denaturation at 95 °C for 10 s, and data collection at 60 °C for 1 min. All reactions were performed with two technical replicates. The expression level was normalised to the level of the reference genes stated and quantified using the ΔΔCt method.

### RNA Fluorescence *in Situ* Hybridisation (FISH)

At the desired timepoint, cells were fixed in 4% (v/v) paraformaldehyde and permeabilised using DPBS containing 0.1% (v/v) Triton X-100 and washed with Stellaris® Wash Buffer A (with 10% (v/v) formamide). Fluorescent probes were then diluted into hybridisation buffer (30 mM saline sodium citrate buffer, 0.2% (w/v) BSA, 10% (w/v) dextran sulphate, 50% (w/v) formamide) to a concentration of 125 nM and 100 μl was placed onto coverslips. Coverslips were incubated in a light-protected humidified chamber at 37 °C for 16 h. Coverslips were transferred to 1 ml Stellaris® Wash Buffer A containing 375 nM Hoechst stain (Sigma) and incubated in a light-protected humidified chamber at 37 °C for 30 min. Coverslips were then washed in Stellaris® Wash Buffer B and mounted for imaging. The cells were imaged using a X60/1.42 NA Oil Plan Apo objective on a DeltaVision Elite system (GE Life Sciences) with an SSI 7-band light-emitting diode for illumination and a monochrome sCMOS camera using SoftWoRks software (version 6). 4’,6-diamidino-2-phenylindole, FITC, TRITC, Cy-5, and DIC channels were used. The images were analysed using the Fiji image processing package (NIH).

## RESULTS

### *In silico* bioinformatic tools predict G-quadruplex formation for the Task3 (GGN)_13_ repeat

5’ Rapid amplification of cDNA ends (5’ RACE) sequencing of full-length Task3 5’ UTR from human brain mRNA revealed a 71 nucleotide (nt) 5’ UTR N-terminal extension, which when sequenced was found to contain a (GGN)_13_ repeat 11 nt downstream of the m^7^G 5’-cap (Figure 1B). GGN repeats can form G4es as well as intermittently bulged duplex secondary structures within mRNA (43). To predict the structures potentially formed by this (GGN)_13_ repeat, we used the *in silico* G4 prediction tools, QGRS mapper (44) and the cG/cC scoring system (45). QGRS mapper designates a score based on the G4 consensus sequence of [G_2+_N_1-7_]_4+_, which can be used in comparison to scores of known G4 forming sequences, such as NRAS G4 (Figure 1C). In contrast, the cG/cC scoring system designates a minimum score of 2.05 for G4 formation. Both methods suggest Task3 (GGN)_13_ is predicted to form a G4 structure when constrained as a (GGN)_13_ repeat (21 for a singular G4 or 59 for 3 tandem G4s) and when in the context of the extended 5’ UTR sequences (Figure 1C). We compared scores for Task3 (GGN)_13_ to the well characterised NRAS 5’ UTR G4 structure (14), a duplex forming control (predicted through RNAfold), and Task3 (GGN)_13_/NRAS G/C matched complementary sequence controls. Task3 (GGN)_13_ and NRAS wild-type (WT) G4 were both predicted to form G4s, whilst the remaining controls were not.

### Task3 (GGN)_13_ forms a parallel G-quadruplex at physiological [K^+^]

To address the sequence prediction scores from Figure 1C, we used circular dichroism spectroscopy (CD) to interrogate the structural conformation of the Task3 5’ UTR (GGN)_13_ repeat using DNA oligonucleotides of various sequences of interest (Figure 1C). DNA oligonucleotides were used in place of RNA due to availability and increased experimental versatility. Given that DNA analogues of parallel RNA G4s show the same G4 forming ability and CD traces, this posed no issues when investigating Task3 (GGN)_13_ G4 formation (46). At a physiological concentration of 150 mM K^+^, Task3 wild-type (WT) (GGN)_13_ forms a parallel G4, exhibiting the spectral maxima (λ_max_) of 210 nm and 261 nm (Figure 1D) previously reported for and consistent with parallel G4 formation (46, 47), which was also observed for NRAS WT G4 (Figure 1E).

To further test G4 formation for Task3 (GGN)_13_, we compared the CD spectra of the WT G4 forming sequence to the complementary (CCN)_13_ sequence to prevent G4 formation without disrupting the ability for any potential duplex formation through canonical Watson-Crick base pairing. At 150 mM K^+^, the Task3 complementary (CCN)_13_ sequence exhibits a λ_max_ at ~280 nm and λ_min_ at 210 nm and 250 nm (Figure 1D), typical of a B-form duplex (48), suggesting the formation of a different, non-G4 structural conformation when compared to Task3 WT (GGN)_13_. From this, it is likely the presence of doublet guanine-tracts that is contributing to the structural conformation observed for Task3 WT (GGN)_13_, and not Watson-Crick base pairing due to the distinct non-B conformation. When complementing the NRAS G4 sequence, a loss of any defining λ_max/min_ is observed, suggesting that no structure is likely to be forming (Figure 1E).

Hoogsteen hydrogen bonding is fundamental to G4 formation. We therefore sought to inhibit G4 formation through pre-incubation of the oligonucleotides with 1% dimethylsulphate (DMS), inducing methylation at the N7 position of guanine and preventing the Hoogsteen hydrogen bonding between N7 and the C2 amino group of neighbouring guanines (49) (Figure S1A). Upon methylation, both Task3 WT (GGN)_13_ and NRAS WT G4 exhibited reduced molar ellipticity ([θ]) at the λ_max_ of 210 nm and 261 nm (Figure 1F and S1B respectively), indicating reduced G4 formation. However, the complementary base sequences and duplex controls showed stabilisation and increase in molar ellipticity at their respective λ_max_ that do not correspond to G4 formation (Figure S1 C, D and E). These data therefore suggest a dependency of Hoogsteen guanine-guanine hydrogen bonding on the structural conformations observed at 150 mM K^+^ for Task3 WT (GGN)_13_ and NRAS WT G4. Further, we observed that Task3 WT (GGN)_13_ G4 formation was independent of oligonucleotide concentration through no change in CD signature of between 1-50 μM oligonucleotides, indicative of an intramolecular G4 as opposed to a multi-strand association dependent on oligonucleotide concentration (data not shown). The G4 structure was maintained when in the presence of the 11 nt upstream and 10 nt downstream of the Task3 5’ UTR (GGN)_13_ repeat, and so was therefore also maintained in a wider sequence context (Figure S2A).

Unlike duplex structures, G4 formation is dependent on the availability of monovalent cations to stabilise the central electronegative pore (4) (Figure 1A). To test the ionic dependency of Task3 (GGN)_13_, we compared G4 formation under different ionic conditions, either K^+^ to permit G4 formation or Li^+^ to inhibit G4 formation. At 150mM K^+^, Task3 (GGN)_13_ forms a parallel G4, exhibiting the CD λ_max_ of 210 and 261 nm (Figure 1G), whilst the duplex control exhibits a λ_min_ of 210 and 250 nm and λ_max_ of 270-280 nm, typical of a B-form duplex (Figure S2B). For Task3 WT (GGN)_13_ we observe a complete shift in spectral shape when folded in the presence of 150 mM Li^+^ (Figure 1G), exhibiting a similar trace to that observed for the duplex control, indicating an absence of G4 formation in Li^+^ and a strong dependency on K^+^ availability for Task3 G4 formation. As expected, the duplex sequence showed no ionic sensitivity, and its CD trace was maintained fully when shifting from K^+^ to Li^+^ (Figure S2B). These results suggest that in the absence of K^+^ and presence of Li^+^, alternative molecular bonding is favoured for Task3 (GGN)_13_ instead of the guanine-guanine Hoogsteen bonding critical for G4 formation, a characteristic typical of G4 structures.

### Task3 (GGN)_13_ G-quadruplex formation is K^+^ dependent

In order to further characterise the K^+^-dependency of G4 formation for Task3 (GGN)_13_, we analysed the CD spectra in the presence of different K^+^ concentrations from 5-150 mM K^+^. As before, we observe 2 distinct structural conformations at low vs high K^+^ concentrations, with spectra indicative of G4 formation only observed at 50 mM K^+^ and above. We observe an increase of approximately 10-fold in molar ellipticity at 210 and 261 nm between 50-150 mM K^+^ (Figure 2A and B). These data suggest a strong dependence on higher physiological K^+^ concentrations for Task3 G4 formation, and also the potential for Task3 (GGN)_13_ to form alternative structures, such as a duplex (Figure 2C) at low [K^+^].

**Figure 2.**
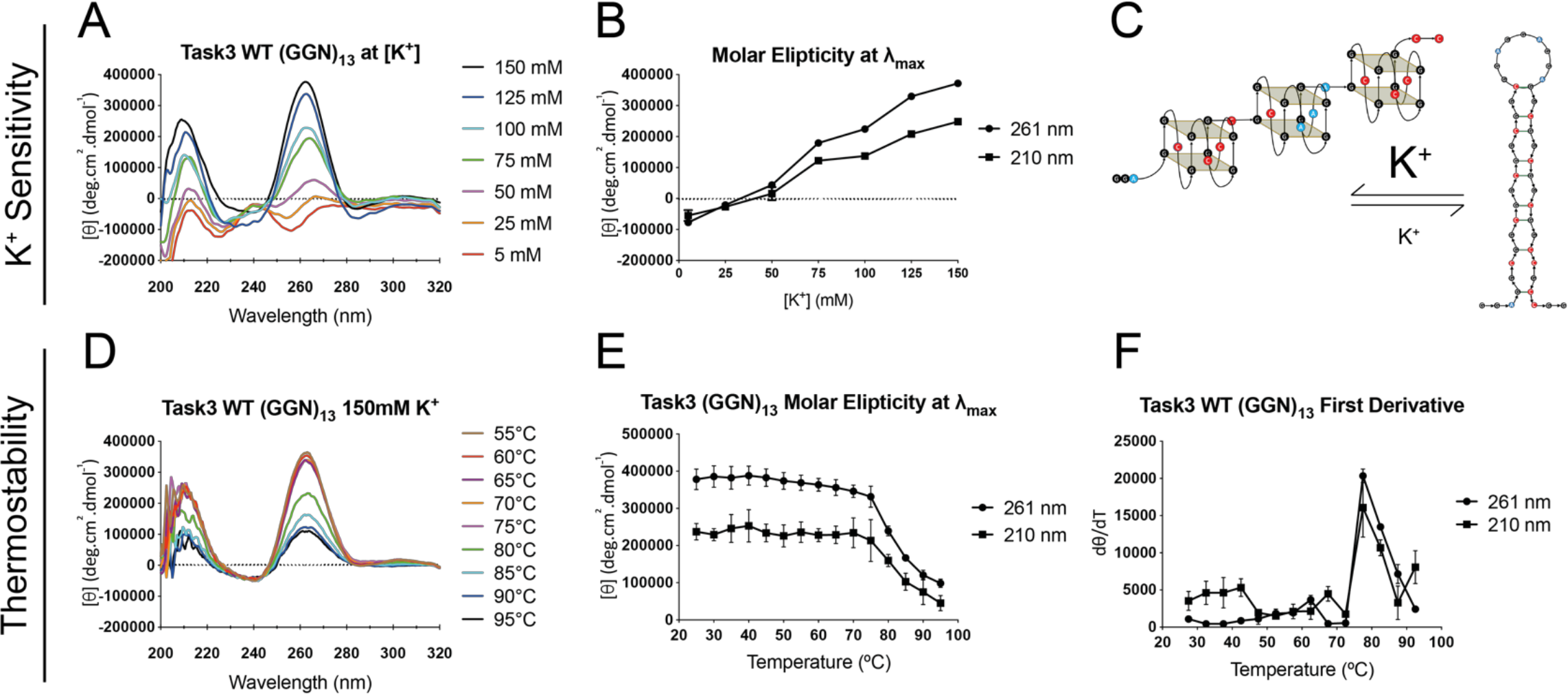
Task3 WT (GGN)_13_ G4 formation is highly thermostable and sensitive to K^+^. (**A**) The CD spectra of 10 μM Task3 WT (GGN)_13_ oligos folded in potassium phosphate buffer and supplemented with KCl to a free [K^+^] of between 5-150 mM. (**B**) Molar ellipticity at the λ_max_ of 210 and 261 nm for Task3 WT (GGN)_13_ at increasing [K^+^]. (**C**) Schematic illustrating the proposed balance of structural conformations of Task3 (GGN)_13_ at low vs high K^+^ concentrations. The CD thermal profile of 10 μM Task3 WT (GGN)_13_ (**D**) which was quantified at the two λ_max_ of 210 and 261 nm (**E**) with estimation of melting temperature through first derivative calculations (**F**). N=3 replicates of 5 repeated acquisitions for signal:noise correction with representative traces shown for each condition.

Conformational shifts from duplexes to G4es have been observed in several instances upon changes in ionic concentrations (50, 51). Here however, we observe a conformational shift only when the ionic composition was discretely changed before folding of the oligonucleotides, with no change in Task3 spectra when Li^+^ was added to oligonucleotides that had been pre-folded in 150 mM K^+^ (Figure S2C), or when more K^+^ was added to oligonucleotides pre-folded in 5 mM K^+^ (Figure S2D). This further indicates that in the presence of low concentrations of K^+^, Task3 WT (GGN)_13_ is not single stranded as increasing K^+^ thereafter does not induce G4 formation and the presence of increased K^+^ is unable to overcome the ordered structure formed at low [K^+^].

### Task3 5’ UTR WT G-quadruplex is thermostable under physiological conditions

G4es are highly thermostable under physiological ionic conditions (3), and so we characterised the thermostability of the Task3 (GGN)_13_ G4 in order to determine whether the G4 melting temperature was above physiological temperature. We performed a CD melt assay using 10 μM experimental oligonucleotides that were folded in 150 mM K^+^ and analysed within a Peltier cooled cell holder from 20-100 °C, with spectra obtained at 5 °C intervals.

Task3 (GGN)_13_ G4 exhibited a highly stable thermal profile, with no reduction in molar ellipticity at the λ_max_ until 75-80 °C (Figure 2D and E). From these data it is evident that the G4 is thermostable under physiological K^+^, with an estimated melting temperature of between 75-80 °C from the first derivative calculation of the melt curve for Task3 WT (GGN)_13_ (Figure 2F). These data suggest that if G4 formation is maintained in the presence of 150 mM K^+^ *in vivo*, the Task3 (GGN)_13_ G4 would remain folded at physiological temperature and could contribute to post-transcriptional regulation of the expression of Task3 channels.

### Task3 5’ UTR G-quadruplex inhibits translation of Task3-FLAG reporter constructs

To study the influence of the 5’ UTR (GGN)_13_ G4 on the translation efficiency of Task3, we generated reporter constructs possessing WT or mutated regions within the 5’ UTR G4 region (Figure 3A) that all produce identical 3× FLAG tagged peptides termed Task3^M1^-3F (Figure 3B). The product of translation consisted only of the N-terminal cytoplasmic and first transmembrane domains in order to avoid formation of functional Task3 K^+^ channels that would alter intracellular K^+^ concentration. Upstream of the coding sequence, the constructs differ only in the G-rich (GGN)_13_ region between positions -120 and -81, with the wild-type containing the 131 nt 5’ UTR, the complementary control containing a (CCN)_13_ repeat mutation within the 39 nt G4 region, and the ΔG4 containing complete deletion of this 39 nt portion, whilst maintaining the rest of the 95 nt 5’ UTR (Figure 3A).

**Figure 3.**
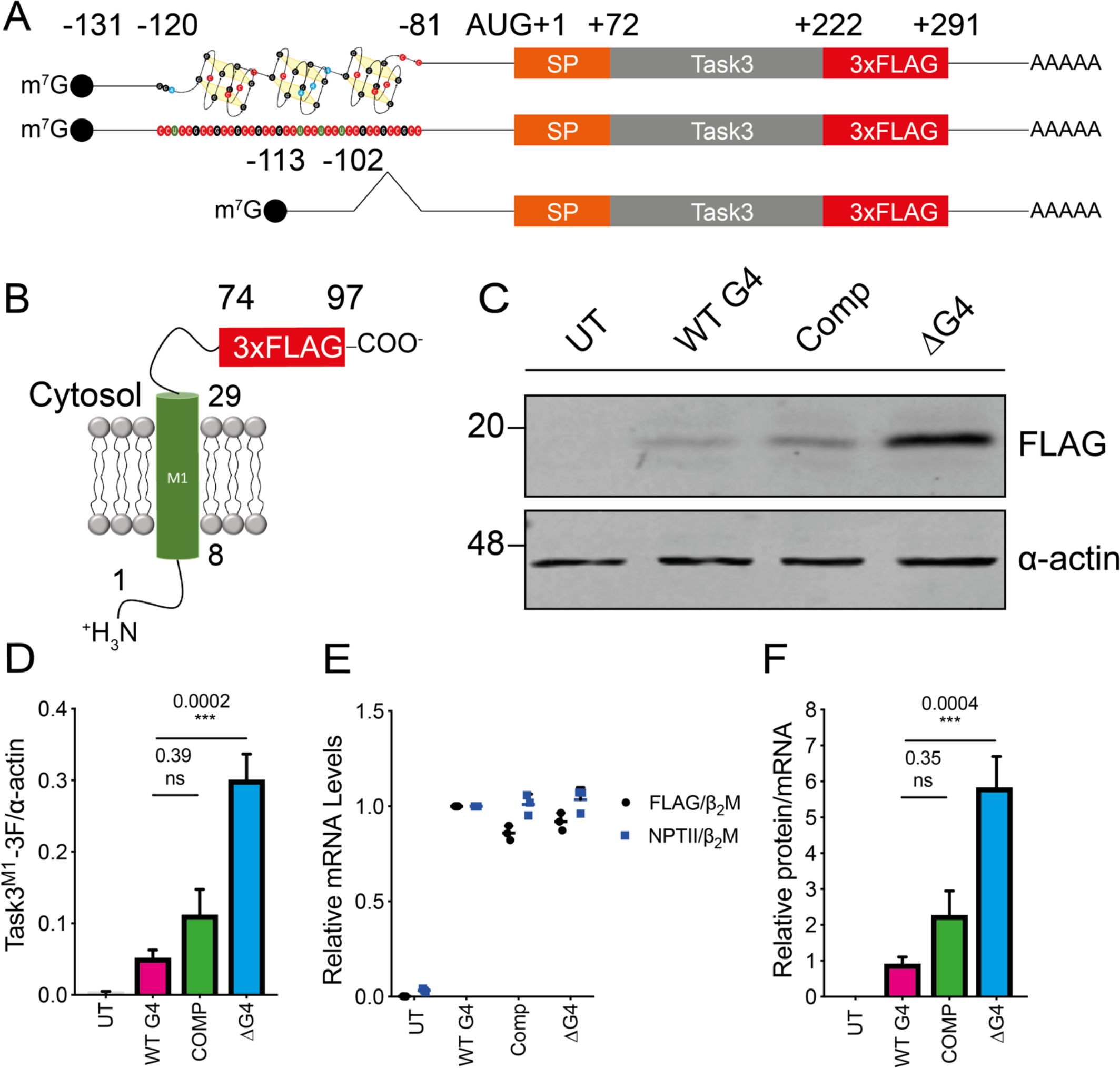
Task3 5’ UTR G4 inhibits translation of Task3-FLAG peptides. (**A**) Schematic of the mRNA transcripts produced by the different Task3 mutant 5’ UTR constructs, possessing either the wild-type full length 5’ UTR, complementary (CCN)_13_ repeat mutant, or a (GGN)_13_ deletion (ΔG4) mutant. SP – signal peptide. (**B**) All constructs translate the identical 97 amino acid peptides (Task3^M1^-3F) allowing for comparison of different translation efficiencies that arise from different structural species present within Task3 5’ UTR. (**C**) Western blotting of HEK-293 lysates transfected with each Task3 mutant 5’ UTR construct. (**D**) Quantification of Task3^M1^-FLAG expression normalised to α-actin expression. (**E**) Mutant 5’ UTR Task3^M1^-FLAG mRNA transcript abundance normalised to endogenous HEK-293 β_2_Μ (Black) or the internal vector control NPTII normalised to HEK-293 β_2_Μ (Blue) for transfection control. (**F**) Relative translation efficiency attributed to each of the wild-type or mutant Task3 5’ UTR mRNA transcripts where Task3 WT G4 5’ UTR = 1. N=3 biological replicates where error bars represent SEM. Statistical analyses were carried out by one-way ANOVA **D** - (F(3, 8) = 25.84, p = 0.0002), **F** - (F(3, 8) = 21.42, p = 0.0004) with post-hoc Sidak’s multiple comparison tests, p-values shown. UT = untransfected.

HEK293 cells were transfected with the FLAG-tagged reporter constructs and protein lysates were analysed through Western blotting 48 hours post-transfection (Figure 3C). Task3^M1^-3F expression was normalised to α-actin and quantified (Figure 3D), and a significant degree of expressional control was elicited in the presence of the 5’ UTR G4. We observed an increase in expression of Task3^M1^-3F by ~6-fold upon deletion of the (GGN)_13_ repeat in the ΔG4 sample, thus suggesting it acts as a key regulatory mechanism of controlling rates of Task3 translation. Interestingly, not all mutants showed the same degree of expression increase, with the complementary (CCN)_13_ repeat eliciting only a partial relief in translation repression compared to the WT G4. This further suggests the formation of an alternate secondary structure such as a duplex or intercalated motif (i-motif), as suggested in Figure 1D. To confirm that the observed increase in Task3^M1^-3F expression was due to G4-induced translational repression as opposed to transcriptional repression, we quantified levels of Task3^M1^-3F mRNA via qPCR (Figure 3E). When normalised to the endogenous HEK293 beta-2-microglobulin (β2M), Task3^M1^-3F mRNA levels remained consistent for all constructs, indicating no changes in levels of transcription. Similarly, levels of neomycin phosphotransferase 2 (NPTII) mRNA, an internal vector control gene expressed from the transfected plasmids, were analysed in comparison to endogenous β2M mRNA to ensure consistent levels of transfection for each construct (Figure 3E). These data suggest that all constructs possess no significant difference in transfection rate nor levels of Task3^M1^-3F transcription.

The data from Figure 3D and E allowed for comparison of relative mRNA to protein levels for Task3^M1^-3F to inform us of the translation efficiency for each construct (Figure 3F). These data further suggest that the increase in Task3^M1^-3F expression upon deletion of the G4 (ΔG4) is due the loss of the translationally-repressive moiety.

### Overexpression of the G-quadruplex helicase DHX36 partially relieves the (GGN)_13_ G4 translation repression of Task3

The DEAH-box helicase 36 (DHX36) is a G4 resolving helicase (52). Binding to G4s increases translation, prevents the accumulation of translationally repressed mRNAs (53) and has also been shown to be important in the neurite localisation of certain RNA species (54). We therefore next investigated whether overexpression of DHX36 in HEK293 cells could relieve the G4 mediated translation repression observed for Task3^M1^ mRNA.

HEK293 cells transfected with a FLAG-DHX36 construct (3F-DHX36) drove an average increase in DHX36 expression of between 10-15-fold, from quantification of total vs FLAG-DHX36 (Figure 4A, left). HEK293 cells were subsequently co-transfected with Task3^M1^-3F WT or mutant 5’ UTR constructs and either FLAG-DHX36 (3F-DHX36) or Firefly Luciferase-FLAG (Fluc-3F) as a co-transfection control. Expression for each condition was normalised to the Task3 ΔG4 to compare translation efficiencies of G4 vs non-G4 mRNA in the presence of DHX36 or Fluc-3F (Figure 4A, right). When co-expressed with Fluc-3F, Task3^M1^-3F maintained a similar expression pattern for the three 5’ UTR variants as was previously observed in Figure 3C-D. However, upon co-transfection with 3F-DHX36, we observed an increase of ~2-fold for the proportion of Task3^M1^-3F WT G4 expression when compared to the ΔG4 control in the presence of 3F-DHX36. In contrast, when the 5’ UTR (GGN)_13_ sequence was mutated to (CCN)_13_, in the presence of DHX36 we observed no significant change in Task3^M1^-3F expression when compared to the Fluc-3F control. These data indicate DHX36 has preferential activity between samples, confirming that Task3 (GGN)_13_ G4 formation is maintained and that the Task3 Comp is forming an alternate structure to a G4, that is still mildly repressive to translation.

**Figure 4.**
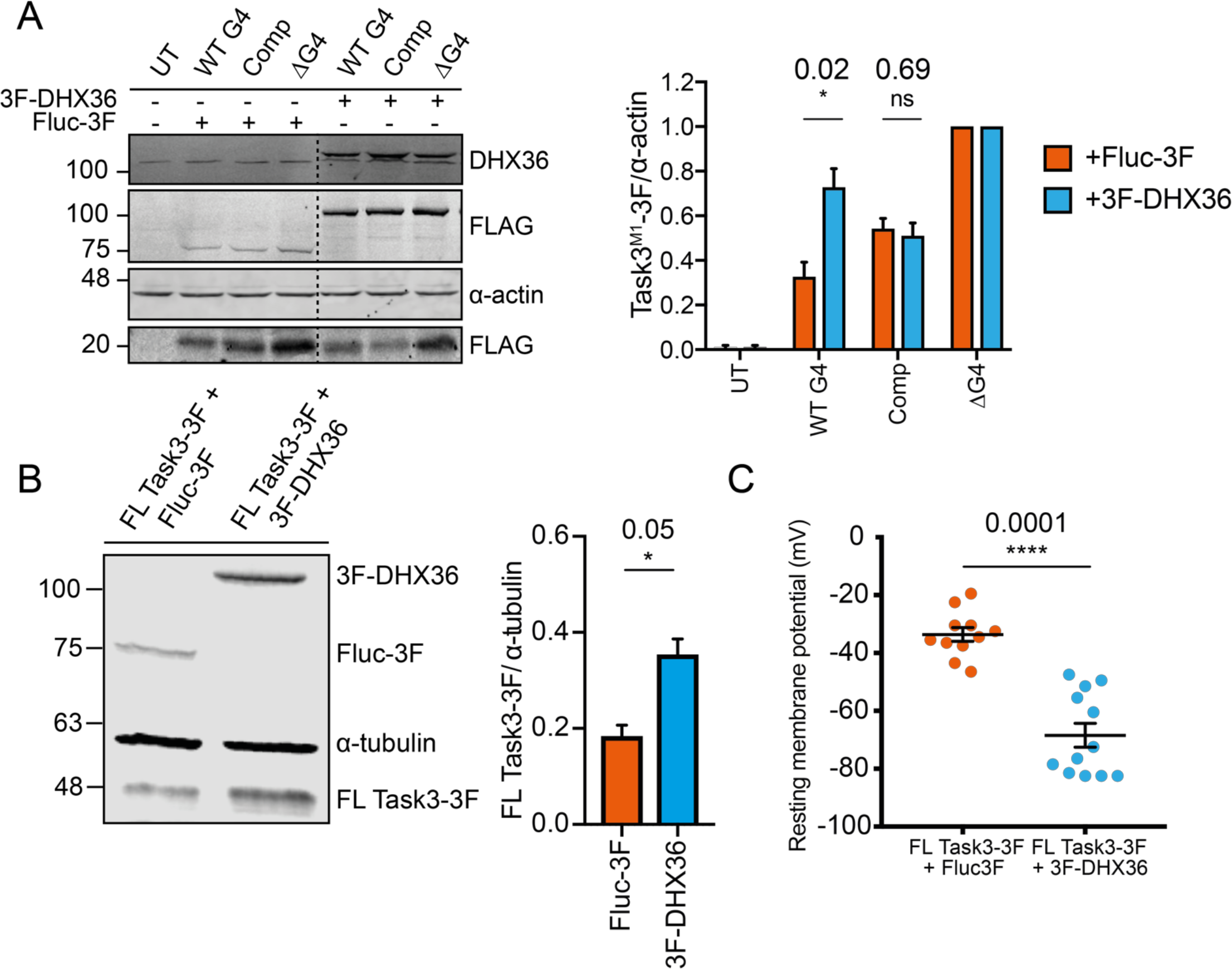
The translation inhibition induced by the Task3 5’ UTR G4 is relieved through overexpression of the G4 specific helicase DHX36. (**A**) Western blotting of HEK-293 lysates co-transfected with each Task3 mutant 5’ UTR construct and either a non-G4 interacting co-transfection control (Fluc-3F) or the G4 resolving helicase DHX36 (Left). Quantification of Task3^M1^-3F expression normalised to α-actin expression. In each co-transfection condition, Task3^M1^-3F expression values for the Task3 5’ UTR mutant constructs were normalised to the ΔG4 control (Right). (**B**) Western blotting of HEK293 lysates transfected with Fluc-3F or 3F-DHX36. Membranes were blotted using anti-FLAG antibody to identify expression of Fluc-3F, 3F-DHX36 as well as FL Task3-3F and anti-α-tubulin for α-tubulin (Left), with exogenous FL Task3-3F expression quantification normalised to α-tubulin (Right). (**C**) Whole cell patch clamp recordings of HEK293 cells co-transfected with FL Task3-3F and either Fluc-3F or 3F-DHX36 (Left). Western blotting of HEK293 cell lysates under the same transfection conditions and time-points (Right). N=3 biological replicates where error bars represent SEM. Statistical analyses were carried out by unpaired one-tailed T-tests, p-values shown. UT = untransfected.

To investigate whether Task3 mRNA was a target of DHX36 endogenously and within the context of its full-length coding sequence, we analysed the expression of endogenous Task3 upon overexpression of DHX36 (Figure S3A). We observe that upon transfection of 3F-DHX36, endogenous Task3 expression was increased by an average of 1.25-fold when compared to the Fluc-3F transfection control. This suggests that Task3 (GGN)_13_ G4 formation is maintained in cells for both endogenous and exogenous transcripts, signified by the preferential relief of translation inhibition of the Task3 WT (GGN)_13_ G4 transcripts from overexpression of the G4-specific helicase DHX36.

### Increasing Task3 expression through DHX36 G-quadruplex unwinding decreases the resting membrane potential of HEK293 cells

Since overexpression of DHX36 leads to an increase in endogenous Task3 expression in HEK293 cells (Figure S3A), we sought to investigate if this overexpression would lead to a Task3 dependent alteration in the membrane potential of HEK293 cells from increased K^+^ leak currents. HEK293 cells display a resting membrane potential of ~40 mV (55). We predicted that an increase in plasma membrane expression of Task3 K^+^ channels would allow us to detect a negative shift in membrane potential. We measured membrane potentials of HEK293 cells transfected with either Fluc-3F or 3F-DHX36, as carried out in previous experiments. When compared to cells transfected with Fluc-3F, we did not observe a significant decrease in resting membrane potential of cells transfected with 3F-DHX36 (−38.75 ± 3.14 mV, −41.25 ± 2.45 mV, p=0.15) (Figure S3B). These data however suggest that DHX36 is unlikely to have a broad spectrum of mRNA targets that would lead to a significant alteration in resting membrane potential, although cells transfected with high concentrations of 3F-DHX36 did show increased cell death (data not shown).

Task3 mRNA abundance is relatively low and our data suggest that the action of DHX36 is likely limited by the amount of available Task3 mRNA for it to resolve. To investigate this, we co-transfected in a full-length Task3 construct possessing the WT 5’ UTR (FL Task3-3F) with either Fluc-3F or 3F-DHX36, in an attempt to increase Task3 mRNA levels and drive a larger decrease in resting membrane potential in the presence of 3F-DHX36 when compared to the Fluc-3F control. Whilst overexpression of DHX36 was able to drive a 1.2-fold increase in endogenous Task3 expression in comparison to FLuc-3F (Figure S3A), an overexpression of both DHX36 and Task3 mRNA was able to further drive an increase in FL Task3-3F expression of 2-fold over endogenous levels (Figure 4B), similar to the level of increase observed for the overexpression of Task^M1^-3F (Figure 3A). This was accompanied by a highly significant decrease in resting membrane potential for FL Task3-3F + 3F-DHX36 compared to FL Task3-3F + Fluc-3F (33.59 ± 2.4 mV, 68.41± 4.14 mV, p=0.0001) (Figure. 4C) suggesting that Task3 channels are incorporated into the plasma membrane, allowing K^+^ permeability of the cell to be increased.

### Distal localisation of Task3 mRNA is dependent on presence of the 5’ UTR G-quadruplex

There is increasing evidence that G4s are involved in regulating local translation of neuronal mRNAs through permitting the localisation of mRNA to these distal compartments (21, 56). We therefore wanted to investigate whether in addition to regulating translation. The 5’ UTR G4 also mediated subcellular localisation of Task3 mRNA. To investigate this, the mutant 5’ UTR Task3^M1^ constructs were used as in previous experiments, but with the FLAG tag substituted for eGFP to provide a longer target sequence for RNA FISH probes, as well as to distinguish between endogenous Task3 mRNA and the experimental G4 exogenous Task3^M1^ transcripts. Subcellular localisation of exogenous WT G4 Task3^M1^-eGFP mRNA was initially determined within murine embryonic cortical neurons through simultaneous FISH and immunofluorescence for the respective axonal and dendritic markers, tau and MAP2 (Figure 5A). Exogenous WT G4 Task3^M1^-eGFP mRNA was localised to both axons and dendrites, where mRNA respectively co-localised with tau and MAP2. We subsequently sought to identify how mutating the 5’ UTR (GGN)_13_ G4 repeat influenced neuronal subcellular localisation of Task3 mRNA to determine whether the Task3 5’ UTR G4 was responsible for neurite localisation.

**Figure 5.**
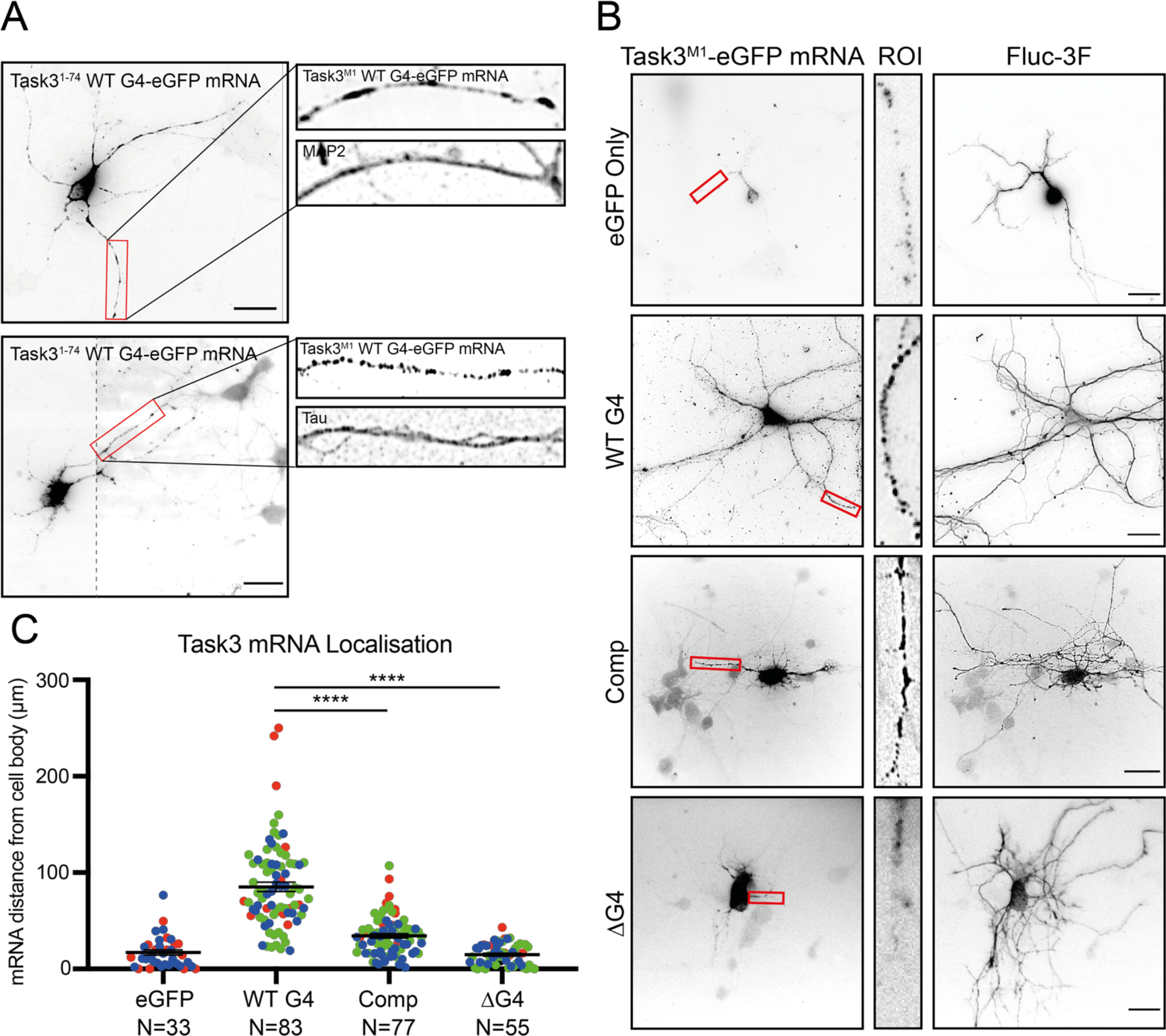
Τask3 5’ UTR (GGN)_13_ repeat is required for distal neurite localisation of Task3 mRNA. (**A**) RNA FISH of DIV10 embryonic murine cortical neurons transfected with WT G4 Task3^M1^-eGFP. Cells were hybridised with anti-eGFP probes and anti-MAP2/Tau primary antibodies to determine neurite localisation of exogenous Task3 mRNA. Image 2 was taken at two exposures to define mRNA granules in both the cell body and neurites with the respective images merged (dotted line). Original un-edited images available in Figure S4. (**B**) RNA FISH of transfected DIV10 embryonic murine cortical neurons co-transfected with mutant G4 5’ UTR Task3^M1^-eGFP and Fluc-3F as a soluble whole-cell tracker. Cells were hybridised with anti-eGFP probes and anti-FLAG M2 primary antibody showing localisation of exogenous mutant G4 5’ UTR Task3^M1^-eGFP mRNA in comparison to whole-cell staining. (**C**) The distance of exogenous mutant G4 5’ UTR Task3^M1^-eGFP mRNA from the cell body, quantified in Fiji for cells possessing neurites ≥ 100 μm in length. Number of neurites analysed is shown from 3 blinded biological replicates (coloured groups), where error bars represent SEM. Statistical analyses were carried out by one-way ANOVA (F(3, 247) = 89.01, p<0.0001), with post-hoc Sidak’s multiple comparison tests; ****p<0.0001. Scale bar = 20 μm.

To address this, the mutant constructs were co-transfected into murine embryonic cortical neurons with Fluc-3F to enable the staining of whole individual transfected cells within the dense neuronal network. Cells were fixed and co-hybridised with anti-eGFP mRNA probes and anti-FLAG M2 primary antibody (Figure 5B). Transfected cells possessing neurites ≥ 100 μm in length were analysed for the distance that exogenous Task3^M1^-eGFP mRNA could be detected from the cell body (Figure 5C).

The Fluc-3F staining indicates that extensive neurite outgrowth was achieved for embryonic murine cortical cells in culture, however, despite consistent levels of neurite outgrowth in each condition, the distance of Task3^M1^-eGFP mRNA reactivity from the cell body of each neuron varied considerably between each of the 5’ UTR G4 region mutants. One-way ANOVA analysis indicates a significant effect of the G4 region manipulation on mRNA distance from the cell body, F(3, 247) = 89.01, p<0.0001. Further post-hoc pairwise comparisons showed that the ΔG4 Task3^M1^-eGFP mRNA showed no significant difference to the eGFP only control, yielding averages of 14.84 ± 1.46 μm and 17.15 ± 2.75 μm respectively (ns, p = 0.99). In contrast, Comp Task3^M1^-eGFP mRNA showed slightly elevated distances for mRNA detection from the cell body in comparison to the eGFP control, with an average of 34.34 ± 2.26 μm (p = 0.02). However, WT G4 Task3^M1^-eGFP mRNA showed the greatest distance of detection from the cell body out of all conditions, with an average of 85.23 ± 4.83 μm (p<0.0001). These data suggest that the presence of either the (GGN)_13_ or (CCN)_13_ repeat within the 5’ UTR was sufficient to drive a significant change in neurite mRNA localisation, with the presence of the (GGN)_13_ G4 forming sequence required for full distal neurite localisation. When comparing within the G4 region mutants, both complementing and deleting the (GGN)_13_ repeat yielded highly significant reductions in distal neurite mRNA localisation when compared to the WT G4, further suggesting that the Task3 WT (GGN)_13_ G4 formation is required for distal Task3 mRNA localisation. Interestingly however, neurite localisation is not completely lost when complementing the (GGN)_13_ repeat as it is when deleting the repeat. As inferred from Figures 1 and 3, it is likely that the (CCN)_13_ repeat forms an alternative secondary structure such as a duplex/stem-loop or intercalated motif (i-motif) due to the CD spectra and translational repression observed. This could also be influencing subcellular localisation of the transcripts through a distinct mechanism to that of G4 driven localisation, potentially mediated by an alternative family of RNA binding proteins (RBP).

### Task3 mRNA is distally localised to neurites in murine embryonic cortical neurons

With the identification of a parallel G4 structure in the 5’ UTR of Task3 mRNA that regulates translational efficiency and localisation of exogenous Task3^M1^ mRNA, we finally wanted to identify how endogenous Task3 mRNA is localised within the context of the full-length endogenous transcripts. To investigate subcellular localisation of endogenous Task3 mRNA in cortical neurons, we used RNA FISH probes targeting mouse Task3 mRNA. Wild type murine embryonic cortical cells were cultured *in vitro* and transfected with eGFP to highlight individual cells within dense neuronal cultures. When tracing neurites from a single eGFP transfected neuron, endogenous Task3 mRNA was observed to be present in both the distal axonal (1) and dendritic (2) projections (Figure 6A).

**Figure 6.**
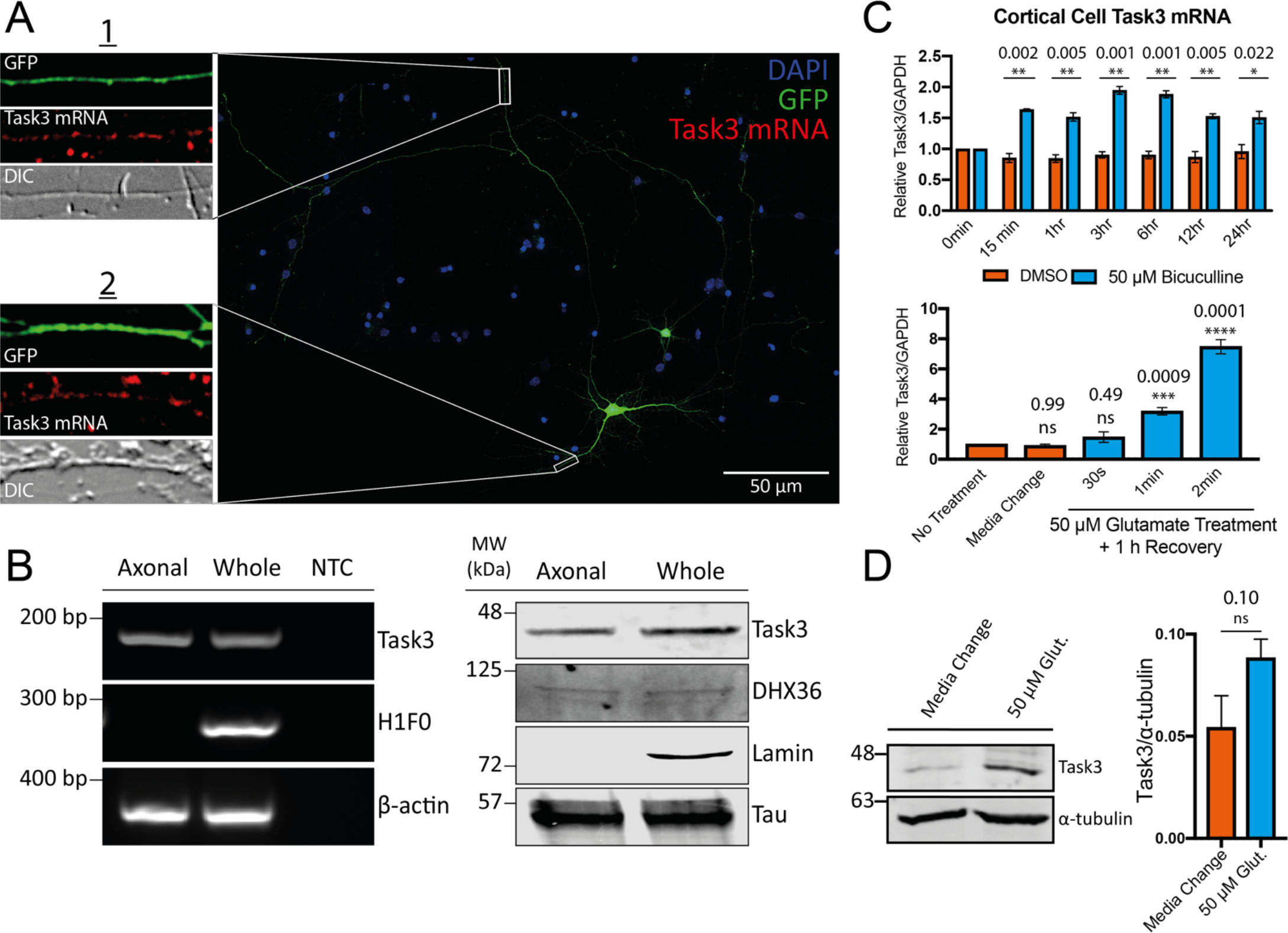
Endogenous Task3 mRNA is localised to distal neurites and upregulates in response to neuronal activity. (**A**) RNA Fluorescence *in situ* Hybridisation (FISH) of eGFP transfected DIV10 embryonic murine cortical neurons hybridised with anti-Task3 mouse probes showing endogenous Task3 mRNA from a singular GFP transfected neuron in distal neurites. (**B**) Primary embryonic cortical neurons were cultured in Corning^®^ Transwell^®^ inserts for the isolation of whole neuron vs axonal mRNA and protein. Western blot (left) and PCR (right) analyses were carried out from whole neuron vs axon protein and mRNA isolates. NTC = no transcript control. (**C**) Total Task3 mRNA levels from DIV10 embryonic cortical neurons treated with 50 μM bicuculline or DMSO over a 24 h period (top), as well as with 50 μM glutamate for 30s, 1min, 2min or a no-treatment control media change with a 1h recovery period (bottom). (**D**) Western blotting of whole cortical cell lysates after 2 minutes treatment with 50 μM glutamate or no-treatment control media change with a 1h recovery period (Left) with quantification of endogenous Task3 expression normalised to α-tubulin (Right). N=3 biological replicates where error bars represent SEM. Statistical analyses were carried out by unpaired one-tailed T-tests or by one-way ANOVA (F(4, 10) = 90.60, p<0.0001) with post-hoc Sidak’s multiple comparison tests, p-values shown.

To confirm the presence of Task3 mRNA and protein in distal axonal projections, cortical cells were cultured in Transwell^®^ inserts (Figure 6B). This allows separation of axonal projections by allowing the growth of a mixed neurite population on the apical membrane face, whilst only an axonal population on the basal membrane face due to constraints in width of the membrane pores (40). RNA and protein extracted from both sides of the Transwell^®^ inserts were analysed through RT-PCR (Figure 6B, left) and Western blotting (Figure 6B, right) for the presence of Task3 and DHX36. The detection of nuclear-localised lamin protein and histone 1 (H1F0) mRNA in only the whole-cell mixed preparation, as well as the detection of Tau protein and β-actin mRNA in both axonal and whole-cell preparations confirms clean isolation of axon-only samples with no cross-membrane contamination. From this, Task3 protein and mRNA were also detected in both preparations, confirming the presence of Task3 mRNA and protein in the distal axon. The presence of Task3 mRNA in both axonal and dendritic projections correlates with that observed for exogenous Task3^M1^ mRNA, suggesting that the G4-mediated control of exogenous mRNA localisation may too be seen for the endogenous Task3 mRNA. Similarly, the presence of endogenous Task3 mRNA distally from the cell body suggests a potential contribution of local translation to Task3 expression.

### Both Task3 mRNA and protein are upregulated in response to neuronal activity

Neuronal activity plays a key role in regulating the spatiotemporal control of local translation that underpins dynamic plasticity at synapses (57–59). Since Task3 mRNA is localised to distal neurites, we sought to examine the role that neuronal activity may play in Task3 mRNA levels, localisation and protein expression. Neuronal cultures contain a finely balanced network of excitatory and inhibitory neurons, with inhibitory neurons becoming responsive at DIV7 and full maturation of network connections at DIV12-13 (60). As a result of this heterogeneous cell population, different pharmacological approaches can be utilised to elicit neuronal activity, with differing intensities and functional outcomes. Bicuculline is a GABA_A_ receptor antagonist that increases neuronal spiking and synaptic protein expression (61), whilst glutamate activates metabotropic and ionotropic receptors including AMPA and NMDA ion channels (62).

We treated wild-type DIV13 murine embryonic cortical neurons with 50 μM bicuculline or a DMSO vehicle control over a 24 h period to induce a low-level increase in basal neuronal spiking through a blockade of inhibitory circuits (Figure 6C, top). Similarly, we also treated DIV13 murine embryonic cortical neurons with 50 μM glutamate for 30 s, 1 and 2 mins with a 1 h recovery period, to induce a short period of intense neuronal activation. RNA was extracted and analysed through RT-qPCR for levels of Task3 mRNA relative to GAPDH (Figure 6C, bottom).

We find that both bicuculline and glutamate are able to induce transcription of Task3 mRNA to differing degrees. Treatment with a DMSO vehicle control induced no change in Task3 mRNA levels, whereas treatment with 50 μM bicuculline induced a modest but consistent increase in Task3 mRNA of between 1.5-2-fold over a 24 h period of stimulation. However, treatment with 50 μM glutamate for just 60s followed by a 1 h recovery period was sufficient to elicit a 3-fold increase in Task3 mRNA, and treatment for 120 s elevated Task3 mRNA levels between 7-8-fold.

In addition to upregulation of mRNA transcription within the nucleus, neuronal activity is also a key driver of regulated local protein synthesis in distal neuronal compartments (63). Given that we observe an increase in levels of Task3 mRNA during activity, we wanted to investigate whether this coupled with an increase in Task3 protein expression that could regulate K^+^ leak currents during periods of prolonged neuronal activity. To analyse this, we treated primary cortical neurons with 50 μM glutamate for 2 mins followed by 1 h recovery, as in the previous experiments for mRNA quantification. Cortical cell lysates were analysed through Western blotting for Task3 expression with 50 μM glutamate compared to a media change control (Figure 6D), where we observe an increase in Task3 protein translation of ~1.6-fold upon treatment with 50 μM glutamate. These data indicate that Task3 mRNA and protein are both responsive to neuronal activity, with Task3 upregulation likely to play a role in maintaining membrane conductance to K^+^ during the neuronal activity changes associated with synaptic plasticity.

## DISCUSSION

*In silico* and bioinformatic analyses predict the presence of over 10,000 potential G4 (pG4) sequences within the human transcriptome (16). Despite recent evidence suggesting that G4s are potentially globally un-folded *in vivo* (64), there is a wealth of evidence in support for G4 formation *in vivo,* ranging from the visualisation of their presence in both the nuclei and cytoplasm of cells through the generation of the G4 specific antibody, BG4 (65), to live cell imaging utilising fluorescent G4-specific ligands such as IMT (66). Similarly, the effect that these G4-specific ligands have on genomic stability as well as transcriptional and translational activity at pG4 regions also suggests maintenance of these structures *in vivo* (11).

In this study, we sought to investigate the functional influence of a 5’ UTR potential G4-forming (GGN)_13_ repeat on the localisation and regulated expression of Task3 mRNA. Through biophysical, biochemical, electrophysiological and cell culture approaches, we have gathered substantial evidence to suggest that this (GGN)_13_ repeat forms a stable parallel G4 structure that inhibits translation of the potassium two-pore domain leak channel Task3 mRNA. Translational repression can be relieved endogenously and exogenously by the primary G4 helicase, DHX36. A shift of the membrane potential (reduction from −33.6 to −68.4 mV) indicates that protein expression and plasma membrane expression also take place and result in modification of K^+^ leak current. Further, our data suggest that the presence of this 5’ UTR G4 is necessary for correct neurite localisation of Task3 mRNA in primary neuronal cultures, providing a new insight into potential mechanisms controlling the regulated expression of K2P channels.

G4s are stabilised by monovalent cations within the central electronegative pore, due to inward orientation of the electron-rich carbonyl groups from each guanine. K^+^ shows the strongest co-ordinating ability due to its ionic radius, with Na^+^ showing weak stabilisation and Li^+^ unable to stabilise G4s (4, 67). K^+^ is the most abundant exchangeable cation in the body, with intracellular concentrations of K^+^ at ~150 mM through the action of different K^+^ transporters/channels (68). We therefore based our biophysical experiments around optimal intracellular K^+^ concentrations of 150 mM. Through CD, Task3 (GGN)_13_ exhibited all of the expected classical G4 characteristics, including a strong dependency on K^+^ concentration, an inhibition of G4 formation through DMS methylation at the N7 position of guanine, and strong thermostability. These characteristics were lost in the absence of K^+^ and also when the sequence was complemented to a (CCN)_13_ repeat, inhibiting G4 formation whilst still allowing any potential GC duplex formation. Similarly, in the absence of K^+^ and when complemented with (CCN)_13_ the CD spectra were similar to that observed for a duplex-forming control. These data suggest that under physiological conditions, Task3 (GGN)_13_ forms a stable parallel G4, however under non-physiological K^+^ conditions, it adopts an alternate structure, likely a duplex in the absence of stabilising K^+^ ions. This was not observed for the NRAS WT G4, where G4 formation was maintained under all conditions except upon DMS-induced N7 methylation of guanine, suggesting that this is likely due to the repeating nature of (GGN)_13_, not just because of G4 formation. *In vivo* this may suggest a more complex equilibrium between the two structures depending on cellular ionic conditions, a process that has previously been observed for other GC rich G4s (50).

When investigating the G4 influence on Task3 expression, several considerations were made to ensure physiological representation. Codon selection and amino acid composition is known to influence translation rate and efficiency post-initiation (69). Similarly, overexpression of K^+^ leak channels perturbs cell signalling and intracellular K^+^ conductance (70), which could therefore influence G4 folding and give a non-physiological representation of the action of the Task3 5’ UTR G4. To overcome this, we designed Task3 constructs that translate the same peptides possessing only the N-terminal cytoplasmic and first transmembrane domains with a C-terminal FLAG tag. This avoided formation of functional Task3 K^+^ channels that would alter intracellular K^+^ concentrations, whilst maintaining physiological codon selection post-translation initiation. We find that the G4 located in the 5’ UTR of Task3 elicited ~6-fold decrease in translation of Task3^M1^-3F peptides compared to when the (GGN)_13_ repeat was deleted, a similar effect to that originally observed for the NRAS proto-oncogene 5’ UTR G4 (14). G4s are prevalent within oncogenes, which has been suggested as a mechanism to buffer the overexpression of oncogenic proteins (71). The overexpression of Task3 and other K^+^ leak channels alters cellular K^+^ conductance and has been implicated in a number of cancers (72, 73), suggesting this 5’ UTR G4 may control the regulated low level expression of Task3 required for normal physiological expression. In contrast, dysfunction and knockdown of Task3 leak channels leads to impaired neuronal migration of cerebral cortical neurons in the developing murine brain and the maternally imprinted intellectual disability Birk-Barel mental retardation (35, 74). Taken together, these data suggest that Task3 expression sits within a very narrow physiological window, which the 5’ UTR G4 may be central to regulating.

DHX36 (also known as RHAU and G4R1) is a G4-specific helicase and has been shown to bind preferentially to G4 and G-rich sequences on over 4500 mRNA transcripts, preventing accumulation of translationally repressed mRNAs (53). Chen et al. recently co-crystallised DHX36 bound to the c-Myc G4, providing valuable insight into the mechanisms underpinning helicase mediated G4 unwinding (75). A mechanism of non-processive unwinding of G4es has been proposed, suggesting a preferential ATP-independent mechanism for DHX36 unwinding of RNA G4s, differing to that for DNA G4s (76, 77). Though Task3 had not previously been identified as a DHX36 target in large scale DHX36 interaction studies, we show here that Task3 mRNA is a functional target of DHX36, thereby relieving the translational repression elicited by the 5’ UTR G4 and increasing cellular K^+^ leak currents specific to Task3. The disparity in changes in resting membrane potential when overexpressing DHX36 in the presence of endogenous vs exogenous levels of Task3 mRNA suggest that Task3 is one of the major targets of DHX36 that would be responsible for increasing K^+^ leak currents. Task3 can be expressed as a homodimer or co-expressed with its paralog Task1 as a functional heterodimer (78), with database sequences for Task1 mRNA suggesting the presence of a similar GGN repeat within its 5’ UTR. Task1 sequencing shows a (GGC)_4_ repeat present at the very 5’ end of the 5’ UTR, which may in fact be an incomplete sequence and might show more similarity in length and composition to Task3 if fully sequenced. Similarly, translation of the ATP-sensitive inward rectifier K^+^ channel KCNJ11 was recently shown to be regulated by a G4 structure within its 3’ UTR (79), which taken together suggest a complex combination of both cis- and trans-acting control mechanisms for expression of Task3 and other potassium channels *in vivo*.

It has been proposed that G4s are also a key driver of subcellular localisation of particular mRNAs. For instance, Subramanian et al., 2011 found that of the identified dendritic mRNAs, approximately 30% had sequences in their 3’ UTRs predicted to form G4 structures (21). Mutating these potential G4 forming sequences showed that delivery of PSD-95 and CaMKIIα mRNA to neurites of cultured primary cortical neurons was dependent on the presence of the G4-forming sequence within their 3’ UTRs (21, 22, 54). Until now, the majority of localisation signals have been attributed to 3’ UTR motifs, however, Muslimov et al. showed that insertion of the FXS associated *FMR1* 5’ UTR (CGG)_24_ repeat into α-tubulin 5’ UTR induced re-localisation from the neuronal somata to dendrites of cultured rat sympathetic neurons (80). Here, we show that the endogenously present (GGN)_13_ repeat, containing two separate GGC tracts of 5 and 4 repeating units, within the 5’ UTR of Task3 mRNA is also able to facilitate neurite localisation. Upon mutation and deletion of the (GGN)_13_ G4 sequence, the distance that Task3 mRNA was detected from the cell body of cultured mouse cortical neurons was reduced by 59 and 83% respectively (Figure 5C), suggesting a dependence on the presence of the (GGN)_13_ G4 structure for Task3 mRNA localisation. Task3 mRNA was also detected as dense puncta along intact neurite projections, as seen through the eGFP and DIC images for both Figures 5 and 6, suggesting that Task3 mRNA is likely to be packaged within messenger ribonucleoprotein (mRNP) complexes, as is observed for distally transported mRNAs within neurons (81–83). Further, the detection of DHX36 in the axonal projections (Figure 6B, right) also suggests the presence of the correct and relevant machinery able to overcome G4-induced translational repression within the same distal compartments as Task3 mRNA.

Current models of G4 regulated local translation of neuronal mRNAs postulate that G4 mRNA is transported in a translationally repressed state by G4 binding proteins (G4-RBPs) such as FMRP. Upon neuronal activity and mGluR activation, G4-RBPs such as FMRP release their G4 containing mRNA thereby partially relieving the translational repression and allowing for spatiotemporally regulated translation of key mRNAs (63, 83). For Task3, we observe both an increase in mRNA concentration and protein translation upon 50 μM glutamate treatment, which could suggest a similar mechanism of activity dependent regulation as is seen for other neuronal G4 containing mRNAs. Axonal transport is reported to occur at speeds of less than 2 μm/s (84). For long-distance neuronal projections, the presence of translationally-repressed pools of mRNA at sites of neuronal activity is key to coupling this rapid increase of protein synthesis, due to delays in anterograde transport from the cell body over long distances. We hypothesise that the increase in membrane expression of Task3 leak channels is a mechanism of maintaining membrane conductance to K^+^ during the neuronal activity. Therefore, a combination of readily-responsive, distally localised mRNA, as well as the slower replenishment of Task3 mRNA to these distal sites would likely be key to facilitating the increase in Task3 protein expression observed in response to neuronal activity.

Taken together, our data suggest that a novel (GGN)_13_ repeat within the 5’ UTR of Task3 mRNA forms a parallel G4 that is responsible for the regulation of localisation and translation of Task3 K2P leak channels. Classical G4 helicases are able to overcome the translational repression elicited by the 5’ UTR G4, thereby increasing cellular K^+^ leak currents and decreasing resting membrane potentials. Task3 is a protein with a narrow window of physiological expression, with over- and under-expression leading to cancer and neurodevelopmental disorders respectively. Given our findings, we suggest that this 5’ UTR G4 is central to the regulated expression of Task3 leak channels *in vivo*.

## DATA AVAILABILITY

5’ RACE sequencing data have been deposited in GenBank [Accession Number: MN510330]

Stellaris® RNA FISH probe designer was used for Task3 mouse Quasar® 570 nm design https://www.biosearchtech.com/support/tools/design-software/stellaris-probe-designer

QGRS Mapper: a web-based server for predicting G4es in nucleotide sequences http://www.bioinformatics.ramapo.edu/QGRS/index.php

## ACKNOWLEDGEMENTS

We thank Professor Keith Fox and Dr Grace Hallinan for critically reading the manuscript.

## SUPPLEMENTARY DATA

**Figure S1.**
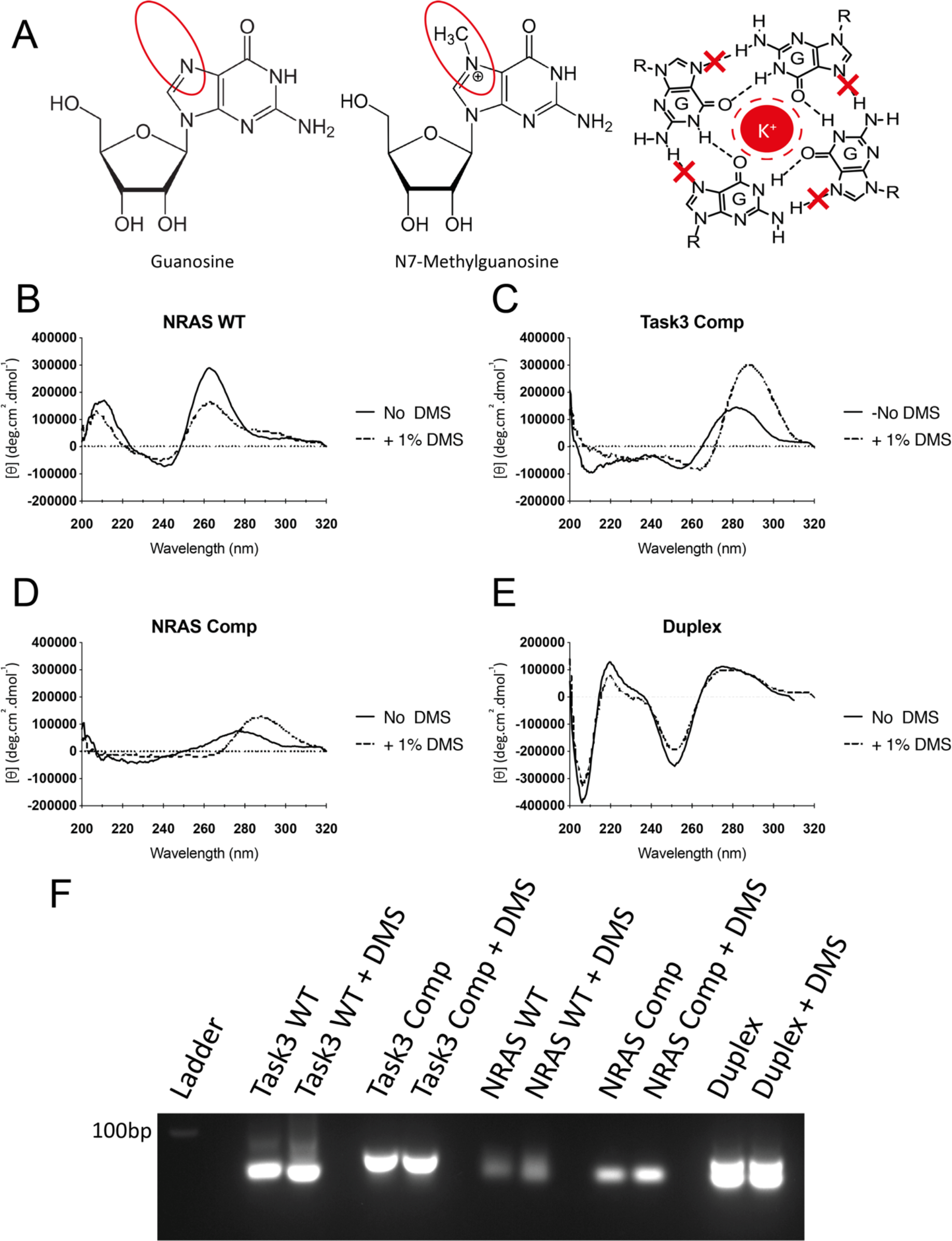
1% DMS treatment inhibits G4 formation for Task3 WT (GGN)_13_ and NRAS WT G4. (**A**) Schematic showing the basis of DMS induced N7-methylation of guanine, inhibiting the Hoogsteen hydrogen bonding underpinning G-quartet formation. (**B**) CD spectra for NRAS WT G4 at 150 mM K^+^ either in the presence of absence of 1% DMS pre-folding. (**C**) CD spectra for Task3 Comp (CCN)_13_ at 150 mM K^+^ either in the presence of absence of 1% DMS pre-folding. (**D**) CD spectra for NRAS Comp at 150 mM K^+^ either in the presence of absence of 1% DMS pre-folding. (**E**) CD spectra for the duplex control at 150 mM K^+^ either in the presence of absence of 1% DMS pre-folding. (**F**) All aforementioned samples run on 1% agarose gel post-CD analysis. CD spectra were obtained from five repeated acquisitions to increase the signal:noise ratio with a representative trace shown.

**Figure S2.**
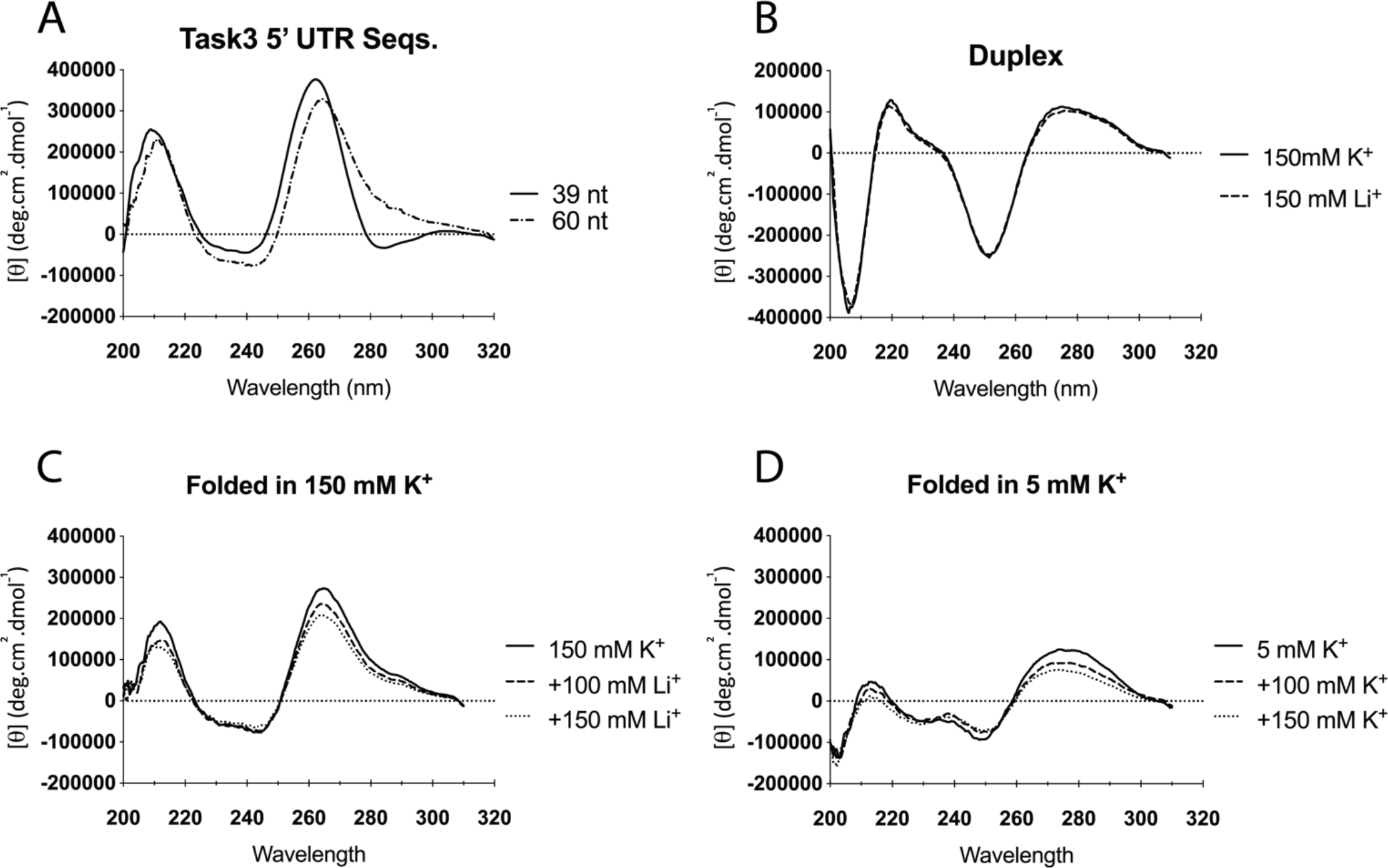
Further characterisation of oligonucleotides via circular dichroism. (**A**) The CD spectra of 10 μM oligos containing either the Task3 WT (GGN)_13_ repeat or a longer 60 nt sequence containing the WT (GGN)_13_ repeat with the 11 nt upstream and 10 nt downstream found within Task3 5’ UTR, folded in potassium phosphate buffer supplemented with KCl to 150 mM K^+^. (**B**) he CD spectra of 10 μM duplex forming oligos folded in potassium phosphate buffer supplemented with KCl to 150 mM K^+^ or folded in sodium phosphate supplemented with LiCl to 150 mM Li^+^. (**C**) The CD spectra for Task3 WT (GGN)_13_ pre-folded in potassium phosphate buffer supplemented with KCl to 150 mM K^+^ and subsequently supplemented with 100- and 150 mM LiCl thereafter. (**D**) The CD spectra for Task3 WT (GGN)_13_ pre-folded in potassium phosphate buffer supplemented with KCl to 5 mM K^+^ and subsequently supplemented with 100- and 150 mM KCl thereafter. CD spectra were obtained from five repeated acquisitions to increase the signal:noise ratio with a representative trace shown.

**Figure S3.**
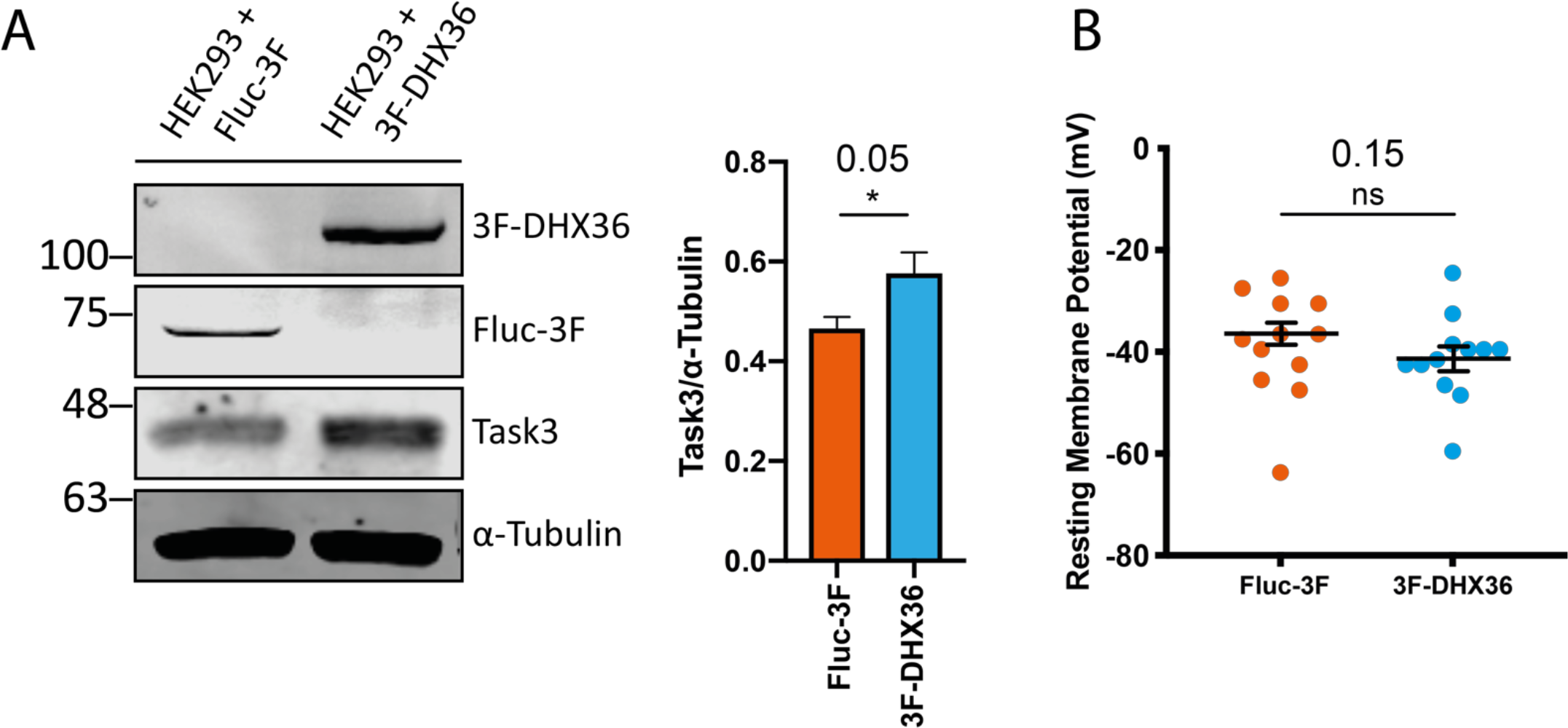
Overexpression of DHX36 increases translation of endogenous Task3. **(A)**Western blotting of HEK293 lysates transfected with Fluc-3F or 3F-DHX36 (Left) and quantified for endogenous Task3 expression (Right). (**B**) Whole cell patch clamp recordings of HEK293 cells transfected with Fluc-3F or 3F-DHX36. N=3 biological replicates where error bars represent SEM. Statistical analyses were carried out by unpaired one-tailed T-tests, p-values shown.

**Figure S4.**
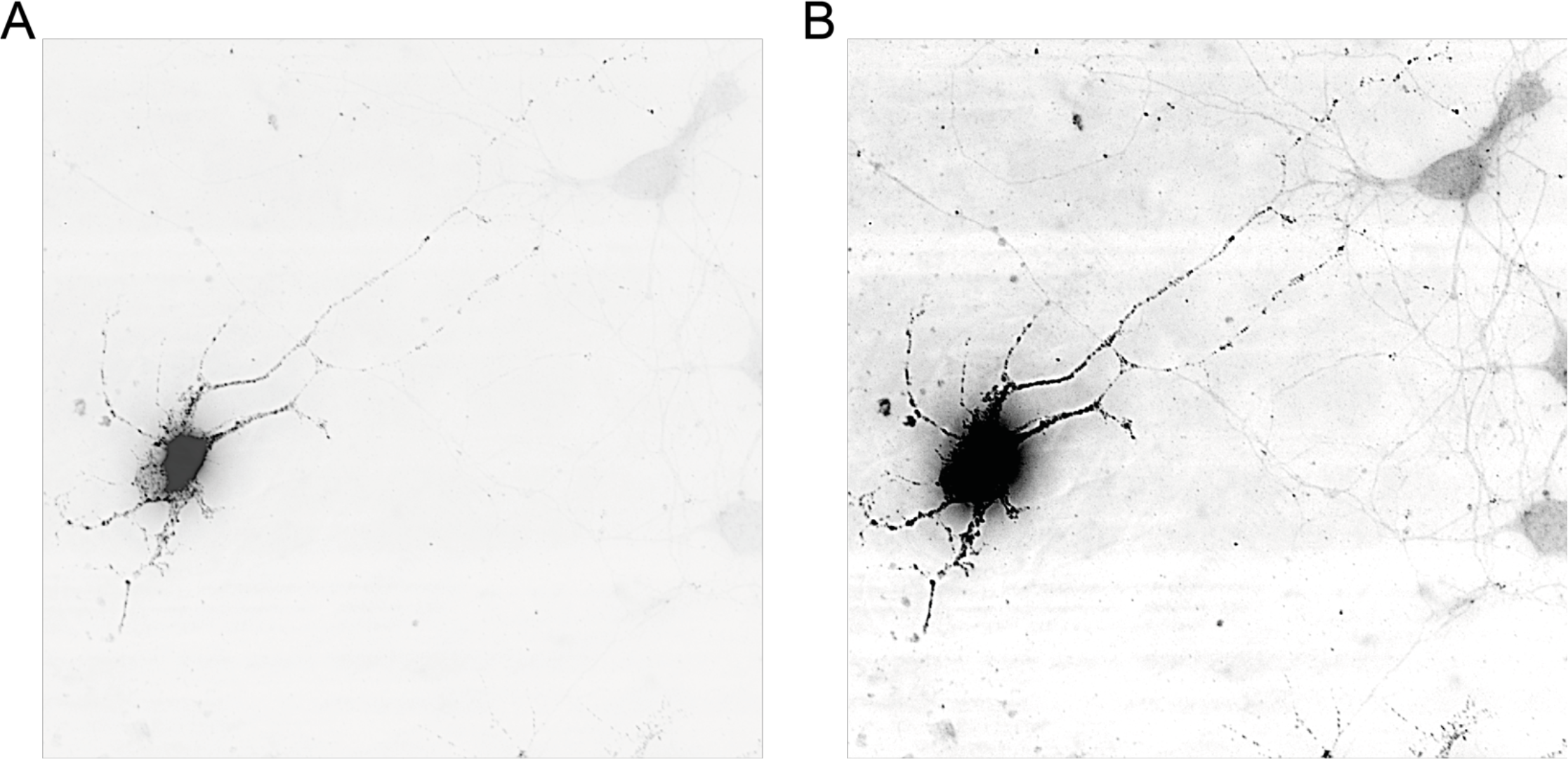
RNA FISH of Task3^M1^-eGFP in DIV10 primary cortical neurons at a short exposure to define mRNA granules within the cell body (**A**), and at a long exposure to define mRNA granules in the neurite projections (**B**).

**Figure S5.**
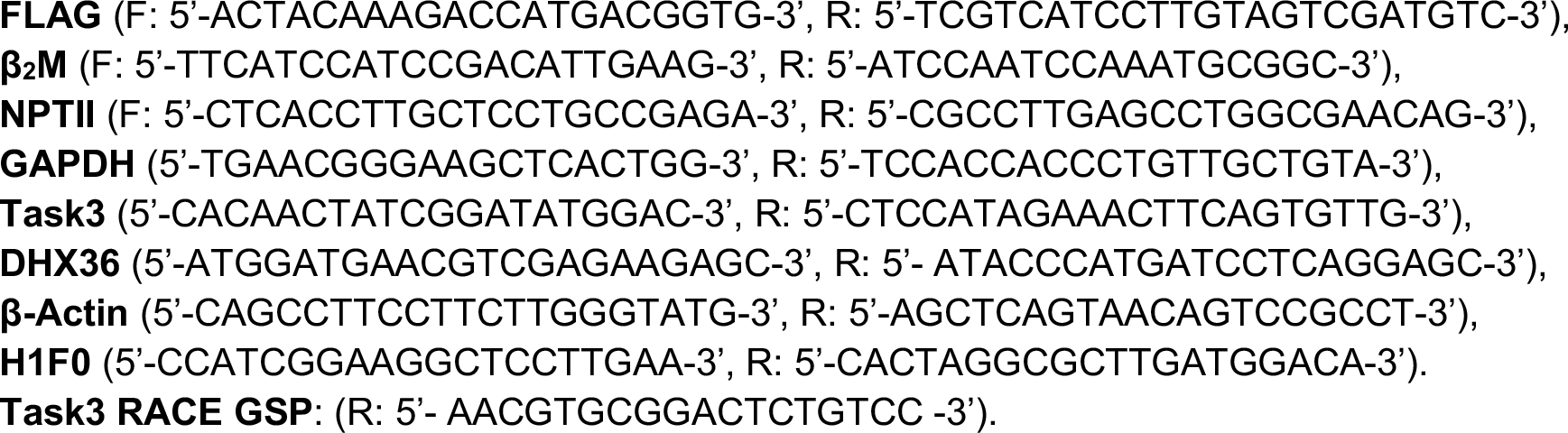
List of primers used for PCR and qPCR obtained from Sigma.

